# The ecological characteristics of the safe sites for early-stage establishment of *Chamaecyparis obtusa* var. *formosana* seedlings in Taiwan

**DOI:** 10.1101/2023.05.15.540728

**Authors:** Shuo Wei, Yu-Pei Tseng, David Zelený

## Abstract

*Chamaecyparis obtusa* var. *formosana* is an ecologically and economically important species in Taiwan, with a high affinity for fog immersion. Our study aims to identify possible stress factors that induced seedling mortality and investigate how different ecological factors influence early-stage safe site requirements of the seedlings. We focused on the effect of large-scale climatic variables, small-scale microhabitat conditions, and biotic interactions on seedling survival and establishment by applying seasonal seedling survival monitoring and establishment survey on both regional and local scale. We identified two alternative ways of seedling death, by environmental-induced mortality and by herbivory. Opposite effects of the same environmental factors on different causes of mortality showed that seedlings might need to balance the risks posed by both causes to optimize their growing conditions. On a regional scale, we observed limited effect of regional climatic variables (namely fog frequency) on seedlings’ establishment and survival but noted a similar seasonal survival pattern among regions. We hypothesize that short-duration droughts during the transition from Plum rain to typhoon season is one of the key mechanisms of environmental-induced mortality. On a local scale, we found that decayed coarse wood debris (CWD) facilitates seedling establishment by providing a “safe site”, likely due to increased colonization of small-stature bryophytes and decreased litterfall accumulation. The effect of bryophytes on seedling establishment varies depending on their thickness, with thicker ones having stronger negative effects. Aside from the bryophytes, the accumulation of litter significantly hindered seedling establishment. We argue that to safeguard the regeneration of *Chamaecyparis obtusa* var. *formosana* population, preserving CWD in the forest floor as a safe site for the seedlings after tree-replacing disturbance in natural forests is essential, particularly under ongoing climate change where more frequent and prolonged drought events are predicted.

**Highlights:**

1. On a local scale, decayed coarse wood debris (CWD) provides “safe site” for the establishment of *Chamaecypairs obtusa* var. *formosana* seedlings.
2. Regional climatic variables had limited effects on seedlings, but all regions had similar seasonal patterns of seedling mortatlity.
3. Facilitation effect of small-stature bryophytes and litterfall avoidance might be the underlying mechanisms behind CWD safe sites.
4. Preserving CWD for seedlings is important in the context of predicted prolong drought events under ongoing climate change.

## Introduction

Subtropical montane cloud forest is a unique and restricted ecosystem characterized by high air humidity, lower temperatures and slow nutrient cycling, and where the unique plant species composition is observed (Bruijnzeel et al., 2011; Helsen et al., 2022). In Taiwan, the subtropical montane cloud forest occurs at elevations from 1400 to 2600 m a.s.l. (Su, 1984; Li et al., 2013). *Chamaecyparis obtusa* var. *formosana* is one of the dominant species distributed in *Chamaecyparis* montane mixed cloud forest typically distributed in upper montane cloud forests with frequent fog immersion (Su, 1984; Li et al., 2013). However, mature individuals of this species were widely exploited in the 20th century for their valuable timber; thus, its natural distribution shrunk considerably (Lee, 1962; Jen, 1995). Considering the current status that the species has reduced and limited distribution, and the fact that it evolved special features to specialize to foggy conditions of cloud forests (Lai et al., 2007; Pariyar et al., 2017), the species might be vulnerable to environmental changes (Lee, 1962; Li et al., 2015). To assure the sustainability of the *Chamaecyparis obtusa* var. *formosana* population, a number of studies have tried to understand the distribution and habitat requirement of the adult trees of the species (Liu, Koh and Yang, 1961; Liu, 1975; Hawk, 1977; Li et al., 2015). However, compared to adult trees, the seedlings’ establishment phase with different responses to the critical environmental factors is the bottleneck that constrains the regeneration of the population (Harper et al., 1961; Grubb, 1977; Harper, 1977).

Since environmental factors critical to the adults might not have the same effects on their seedlings, it is important to distinguish the environmental requirements of the seedlings from their conspecific adult trees. The seedlings require a *safe site* (Harper, 1977) that provides essential elements but also minimizes hazards to optimize individual growth and establishment (Harper, 1977; Grubb, 1977). When characterizing the properties of a safe site, Harper (1977) highlighted the role of abiotic filters, such as light, water, nutrient, temperature, or substrate conditions, and biotic filters, such as pathogens, herbivores, competitors, or facilitators. These factors filter out the seedlings at each development stage, determining their final fate (Moles and Westoby 2004; Larson and Funk, 2016). In addition, studies have found that seedlings of other small-seed late-successional conifer species require unique microhabitats as safe sites for successful establishment (Simard et al., 1998; Cornett et al., 2000; Mori et al., 2004). In particular, the association between coarse wood debris (CWD) and successful establishment has been observed across different geographical regions for species of *Picea* spp. and *Tsuga* spp. (Harmon et al., 1986; Harmon and Franklin, 1989; Ježek, 2004).

*Chamaecyparis obtusa* var. *formosana* has been reported to regenerate on the CWD under small gaps after tree-replacing events (Liu, 1975; Hawk, 1977; Liao et al., 2003a, b). Small gaps on top of CWD have low-to-moderate light intensity and exhibit high light dynamics caused by sunflecks (Lai et al., 2005). Such light regime is likely aligned with the seedlings’ physiological characteristics, since they can sustain their growth rate by responding rapidly to sunflecks by investing more resources in expanding leaf area (Lai et al., 2005). Aside from the light conditions, limiting their growth to CWD might result from the need to escape the thick litter layer and dense coverage of understory vegetation (Liu, 1975; Hawk, 1977; Chang et al., 2001; Liao et al., 2003a). CWD is also often associated with various bryophyte carpets, which were suggested to either provide better water or nutrient regime, or compete with seedlings for resources and interfere with their emergence. In Taiwan, the facilitation effect of bryophyte carpet colonizing the top of the CWD was reported on a swampy site (Liao et al., 2003b), but its effect on seedlings establishment was not conclusive (Su et al., 2018). On the other hand, in silvicultural literature, *Chamaecyparis obtusa* var. *formosana* has been reported to establish on mineral soils in exposed sites after natural disturbances or clear-cutting (Matuura, 1942; Chang, 1961, 1963; Kuo, 1995), even though such habitat might not be the optimal safe site for the seedlings, since they are less adapted to full-light conditions due to their morphological and physiological characteristics (Matuura, 1942; Zobel et al., 1978; Zobel and Liu, 1980; Lai et al., 2005). Despite a number of studies focused on the possible regeneration pattern of *Chamaecyparis obtusa* var. *formosana*, to our knowledge, no study has identified what stress factors may induce seedling mortality during the early-stage establishment phase. Lacking knowledge of the stress factors on the seedling establishment might lead to overlooking the critical environmental factors forming the safe site, and also limit our ability of understanding the population dynamic.

Under ongoing climate change, several studies have predicted an uplift of cloud bands in future climate scenarios, causing changes in temperature and precipitation patterns that might represent significant stress for *Chamaecyparis obtusa* var. *formosana* seedlings (Still et al., 1999; Hu et al., 2016). Therefore, it is important to identify the stress factors and understand how the seedlings respond to these stresses under varying regional and local environmental conditions. Our study aims to enhance the current understanding of the early-stage safe site characteristics of *Chamaecyparis obtusa* var. *formosana* seedlings by focusing on two main aims. Firstly, we attempt to identify the stress factors that cause the death of the seedlings during their early-stage phase. Secondly, we quantify how regional climatic variables, local biotic and abiotic factors, and their interaction influence the establishment and survival of *Chamaecyparis obtusa* var. *formosana* seedlings across a fog frequency gradient in their natural distribution range. Improving the knowledge of the safe site requirements of *Chamaecyparis obtusa* var. *formosana* seedlings, and better understanding their population dynamics, will provide a theoretical base for the development of effective management strategies for natural forests in Taiwan.

## Materials and Methods

### 1. Study area

The study was carried out across the northern distribution range of *Chamaecyparis obtusa* var. *formosana* in the Xueshan mountain range in Taiwan, between 1700 to 2200 m a.s.l. The study region spans from 121.445° E to 121.365° E and from 24.727° N to 24.524° (Fig. 1; Table 1). Most of the soils belong to Entisols, Inceptisols and Spodosols, which have low fertility, low pH, shallow soil depth, and high organic carbon content (Chen et al., 2015). The study region is located in the ever-wet northeast-inland climatic region, characterized by high annual precipitation (exceeds 2000 mm.) and a high ratio of winter precipitation from the winter monsoon, without significant dry season (Su, 1985). Mean temperature in the warmest month is 13.5 ℃, ranging from 12.5 to 14.0 ℃. The warmest mean monthly temperature is 19.5℃ in July, while the coldest is 6.4℃ in January (Lin et al., 2018). The forests within our study region are represented mainly by subtropical *Chamaecyparis* montane mixed cloud forest vegetation type (Li et al., 2013; Li et al., 2015). The forest physiognomy changes along the gradient of decreasing fog frequency from the east to the west. In the foggiest region in the eastern part, the forests are dominated by *Chamaecyparis obtusa* var. *formosana* with boggy soils and high bryophyte coverage on both trunks and root swallows. As the fogginess decreases toward the west, the forests shift to be dominated by both *C. obtusa* var. *formosana* and *Tsuga chinensis* var. *formosana* in the canopy with *Yushania niitakayamensis* or *Plagiogyria formosana* dominated in the understory (Liu 1975; Li et al., 2013; Li et al., 2015). Most areas are natural forests without any previous management or without logging activity (including selection cutting and CWD harvesting) for at least 30 years. In our study, we avoided localities with recent stand-level cutting or natural disturbances.

**Fig. 1.**
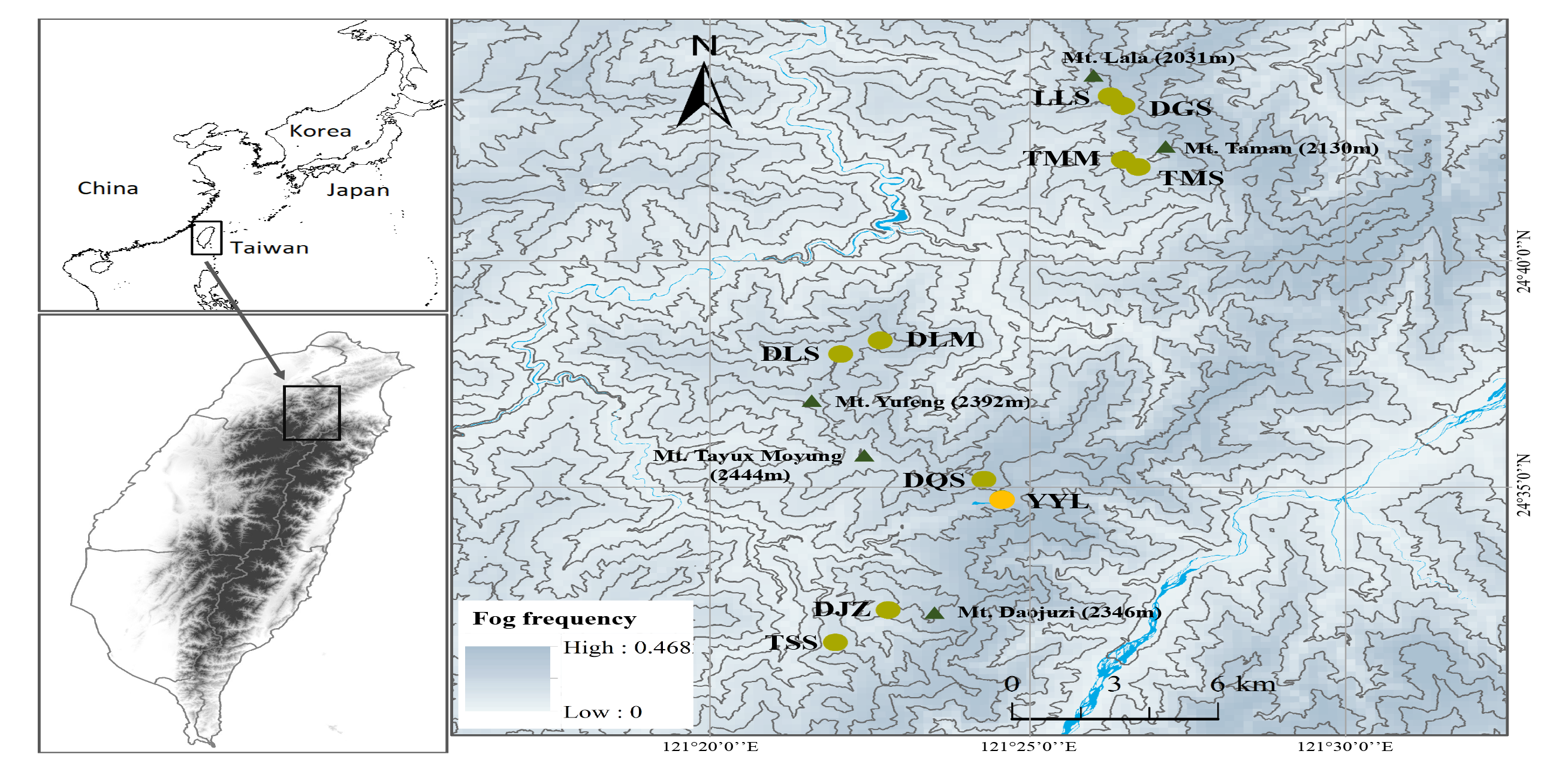
The map of selected sampling regions along a fog frequency gradient in northern Taiwan, located in eastern Asia (abbreviations see Table 1). The yellow-green circles were selected from both seedling survival monitoring and establishment survey. Yellow circle in Yuan-Yang Lake (YYL) was the additional region for survival monitoring. Blue hue represents the fog frequency derived from Schulz et al. (2017) and grey lines is the elevation.

**Table 1.** Geographical coordinates, elevation and regional name (with abbreviation) of the sampling regions.

| Region | Abbreviation | Latitude | Longitude | Elevation |
| --- | --- | --- | --- | --- |
| Donqiusishan (東丘斯山) | DQS | 24.5858° | 121.4041° | 1921 m |
| Lalashan (拉拉山) | LLS | 24.7275° | 121.4377° | 1910 m |
| Daguanshan (達觀山) | DGS | 24.7242° | 121.4403° | 1764 m |
| Dilushan summit (低陸山頂) | DLS | 24.6317° | 121.3666° | 2134 m |
| Dilushan mountainside (低陸山腰) | DLM | 24.6364° | 121.3784° | 1994 m |
| Daojuzishan (島俱子山) | DJZ | 24.5375° | 121.3795° | 2004 m |
| Three Sisters (三姐妹) | TSS | 24.5245° | 121.3652° | 2040 m |
| Tamanshan summit (塔曼山頂) | TMS | 24.7007° | 121.4447° | 2051 m |
| Tamanshan mountainside (塔曼山腰) | TMM | 24.7042° | 121.4412° | 2000 m |
| Yuanyang lake (鴛鴦湖) | YYL | 24.5782° | 121.4095° | 1717 m |

### 2. Sampling design

#### Seedling establishment survey

To examine whether the relationship of *Chamaecyparis* seedlings establishment on regional-scale climatic variables and local-scale microhabitat properties changes along the fog frequency gradient, we designed observational study in localities stratified along fog frequency gradient. We stratified the northern distribution range of the species in Xueshan mountain range along remote-sensed ground fog frequency map obtained from Schulz et al. (2017) into seven levels, and removed the lowest level since it was out of the distribution of the studied species. Several candidate regions were visited to identify localities with forests dominated by *C. obtusa* var. *formosana* in the canopy that are without natural or human disturbances and have feasible accessibility. Finally, nine sampling regions were identified for the following sampling (Fig. 1).

Since the seedlings are sparsely distributed in the forests, a case-control study design, which is frequently used in epidemiology to uncover the causes of rare diseases (Mann, 2003), was preferred over random sampling. This design includes a set of case plots containing studied objects and control plots, specifically selected not to contain the studied object. Within each sampling region, we started from a random position and searched for a plot containing seedlings (further called the *case plot*) with the criteria of having at least five seedlings (excluding first-year seedlings and seedlings taller than 15 cm) in a 0.25 m^2^ area. Once the case plot was found, four plots not containing seedlings (the *control plots*) were located around it to capture the high variability of microhabitat properties and increase the power of the statistical analysis (Woodward, 2013).

We chose the first control plot in a random aspect and random distance (between 5 to 8 m) from the case plot; each next was chosen in an aspect 90° more than the previous one, again in a random distance (5–8 m) from the case plot. The distance was set since we assume the microclimate and seeds availability were similar in this range. Also, we assured each case plot was at least 20 m apart from another case plot, and each control plot was at least 5 m apart from another one. If the location chosen for the control plot had a seedling occurrence or was located on a trail, on a living tree with a basal diameter larger than 20 cm, or inaccessible topography, we extended the distance 2 m ahead to locate the control plot. All the plots were sampled in the summer (from late June to early August) of 2022. We measured a set of abiotic and biotic factors at plot level to quantify the microhabitat properties and estimate the intensity of species interactions. Each plot was placed on a homogenous substrate that was categorized into the following types: soil, living tree, organic matter mat, and coarse wood debris (CWD) (see Table A.1 for detailed description). CWD was divided into heavily-decayed CWD, moderately-decayed CWD, and non-decayed CWD according to their decay status following the 7-class classification system (Lee et al., 1997). Organic matter mat (mat in short) referred to the layer of organic matter accumulated on top of non-decayed CWD or living root swallow with a depth of 5–10 cm. For substrate types other than soil, distance above the soil surface was measured. Bryophyte and litter cover were estimated visually on the scale with 1 % increment if the cover was lower than 10 %, and the scale with 5 % increment if the cover was higher than 10 %. The thickness of both bryophyte and litter cover were estimated twice in each plot with 0.5 cm accuracy using a penetrometer. Canopy cover was estimated visually by accounting for the sphere of 120 degrees above the plot to the nearest 5 % to capture the incident radiation of the understory seedling light regime (Canham et al., 1990). As for biotic factors, we estimated cover of herbs, seedlings (height less than 50 cm), and saplings (height between 50 and 200 cm), using Braun-Blanquet combined abundance-dominance scale (Westhoff and van der Maarel, 1978).

**Table A.1.** Key characteristics of substrate type determination. Coarse wood debris (CWD) decay class criteria were adapted from the 7-class classification system of Lee et al. (1997).

| Substrate type | Decay class | Depth (cm) | Description |
| --- | --- | --- | --- |
| None-decayed CWD | 1–2 | $\leq 1$ | Log or stump sound or undecayed; twigs mostly lacking or present; < 50% bark missing |
| Moderately-decayed CWD | 3–4 | 1–9 | 50–100% bark missing; no to some branch remaining; wood soft with small crevices and small pieces lost |
| Heavily-decayed CWD | 5–7 | $\geq 10$ | >50% of large wood fragment remain, outline of trunk deformed and well decayed |
| Organic matter mat | - | 5–10 | The layer of organic matter accumulated on top of non-decayed CWD or living root swallow |
| Living tree | - | 0 | Living root swallow or trunk of standing live trees without organic matter accumulation |
| Forest soil | - | vary | Substrate type which could not be categorized into aforementioned substrate types. |

#### Seedling survival monitoring

To monitor the seedlings survival in different substrate types across different sampling regions with various fog frequencies, we subjectively set up four 1 m × 1 m plots in each of the study regions selected for seedling establishment survey, which were at least 20 m apart from each other. The plots were set in the summer of 2021, a year before the seedling establishment survey, but in the same localities with an additional sampling region beside Yuanyang lake (Fig. 1). Each seedling of *Chamaecyparis obtusa* var. *formosana* within the plot was numbered with a 6-cm-high plastic tag, which was plugged in the substrate beside the seedling. For four seasons, namely in December 2021 (autumn), March 2022 (winter), June 2022 (spring), and September 2022 (summer), we resurveyed all the plots to check whether the tagged seedlings were alive.

We recorded the individual height and the environmental conditions at the individual-and plot-level. Individual-level factors included whether the individual was established on bryophytes, litter, or substrate itself, the distance of the individual to the soil surface, and whether it was covered by litterfall (litter covered). All plot-level factors mentioned in the previous section were also included and measured by the same methods. The withered seedlings were recorded to be died from environmental-induced mortality. On the other hand, during the resurvey, we observed that some seedlings in the plots have been clipped with obvious clear-cutting edge, resulting in a loss of biomass, or went completely missing in groups with broken plastic tags that did not seem to result from environmental factors. We suspected this phenomenon resulted from the browsing by the herbivores; therefore, we additionally recorded these individuals as being browsed. If the seedlings were killed by browsing, they would be recorded as being died from herbivory. Other seedlings that could not be found during the resurvey were marked as missing.

### 3. Statistical analysis

#### Seedling establishment survey

We used conditional logistic regression to condition on the probability of being a case (non-random sampling from the population in case-control design) (Breslow and Day, 1980). Furthermore, since we sampled four to five 1:4 match sets of case-control plots within each sampling region, we applied mixed-effects matched conditional logistic regression to accommodate the nested data structure (Revelt and Train, 1998; Keating and Cherry, 2004). The logarithm regression coefficient should be interpreted as odds ratio (OR).

All analyses in this study were conducted in R (R Core Team 2022). Owing to our sampling design, sampling regions were regarded as a random effect and each match set was treated as “stratum” in which the plots within each stratum were matched in the analysis. The ‘mclogit’ function in the *mclogit* package and “clogit” function in *survival* package (Elff, 2022; Therneau, 2022) were used. We built a full model that included substrate types, herb, seedling and sapling cover, bryophyte and litter thickness/ cover, and canopy cover as explanatory variables. We applied a stepwise model selection procedure to obtain the most parsimonies model with the selection criterium based on Akaike’s Information Criteria (AIC) and likelihood ratio (LR) test to find the final model. Polynomial terms of each selected variables were examined by LR test separately.

Multicollinearity between explanatory variables of the final model was inspected using generalized variance inflation factor (Fox and Monette, 1992). No serious multicollinearity was detected since no variable exceeded the value of 5. The final model was evaluated with adjusted McFadden’s pseudo-R^2^ (McFadden, 1974) and concordance (Harrell, 2001).

To investigate the effect of regional climatic variables on the seedling establishment, we added the interaction between mean fog frequency or precipitation and substrate types. The interaction term was built to examine the interplay between microhabitat properties and regional climatic variables. The model extended for climatic variable was compared to the final model using the LR test.

#### Seedling survival monitoring

In our data, the seedlings could die either as a result of environmental-induced mortality or by herbivory; therefore, we considered there were competing risks in our data. To accommodate this situation and statistically analyze the explanatory variables’ relationship to the seedlings’ survival time, we applied cause-specific Cox proportional hazard (PH) models that separately modeled the death of seedlings resulting from either environmental-induced mortality or herbivory (Cox, 1972; Lau et al., 2009). The model estimated the instantaneous risk of death for each individual. Survival time was calculated as the number of seasons between the first survey and the seedlings marked as dead. Seedlings that were still alive at the final survey season were coded as right-censored, i.e., the survey time is incomplete. Those seedlings that were missing were also coded as censored. Those seedlings who died from the other competing risk would be treated as censored during the analysis. We used the cumulative incidence function with pointwise confidence interval to visualize death events over time under competing risks (Allignol et al., 2011).

For the survival analysis, we first examined the effect of regional climatic variables on the seedlings’ survival. We applied mixed effect Cox PH model using *coxme* package (Therneau, 2022) with plot nested in region. We included mean seasonal fog frequency, precipitation, and temperature as targeted climatic variables and modelled the effect as time-dependent or non-time-dependent variable. Time-dependent variable has its unique value at each time step (season), while non-time-dependent variable has fixed value that does not change through time representing regional climatic characteristics.

We further explored which individual-and plot-levels factors were associated with seedling survival. We constructed the full Cox PH model containing individual-level (seedling height, distance to the soil surface, located on bryophyte, litter, or substrate, litter covered, being browsed) and plot-level (herb, seedling, and sapling cover, density of the seedling, bryophyte cover and thickness, litter cover and thickness, canopy cover, and substrate types) variables as explanatory variables using “coxph” function in *survival* package for each cause-specific Cox PH model, (Therneau, 2022). Seedlings that were being browsed and being covered by litterfall were not included in the analysis of the herbivory-specific Cox PH model. Then, a stepwise model selection procedure was applied with the selection criteria based on AIC. We found some selected variables did not satisfy the proportional hazard (PH) assumption. To fulfill the assumption, we added the time-varying variable to model the effect of the variables changed linearly through time.

Since our data were collected every season, the data had strong ties; therefore, we used the Effron approximation instead of the Breslow approximation (Hertz-Picciotto and Rockhil, 1997). Also, the “coxph” function could not handle a two-level nested structure of the data. To accommodate the data structure, we used a stratified Cox PH model with a robust standard error that stratified the Cox model by regions and estimated the robust standard error to account for the within-plot nested structure (Barlow, 1994; O’Quigley and Stare, 2002). Multicollinearity between explanatory variables of the final model was checked using the variance inflation factor. No variable exceeded the value of 5, indicating no serious multicollinearity. All explanatory variables, except for the categorical ones, were standardized to zero mean and unit variance before the analysis. Pseudo-R^2^ and concordance of the model were also reported (O’Quigley et al., 2005).

## Results

### Seedling establishment survey

In each of the nine sampling regions, eight sets of plots (each containing one case and four control plots) were sampled; in the Tamanshan summit region, additional two plot sets were sampled. This results in a total of 370 plots from 74 plot sets. In each case plot, an average of 9 seedlings were found (a range between 5 and 38 seedlings). For the substrate types (Fig. 2), soil was the most common substrate type occurring in 65% of all plots (n = 239). Each of the remaining substrate types occurred in less than 10% of plots. When considering only case plots, more than half were found on heavily-decayed CWD (26 case plots, i.e. 35% of all case plots), and moderately-decayed CWD (24, i.e., 32%). Mat (9 case plots), non-decayed CWD (7) has low seedling occurrence. Only one case plot was found on a living tree. Even though the soil was the most prevailing substrate type among all plots, only seven of them had seedling occurrence (7 case plots).

**Fig. 2.**
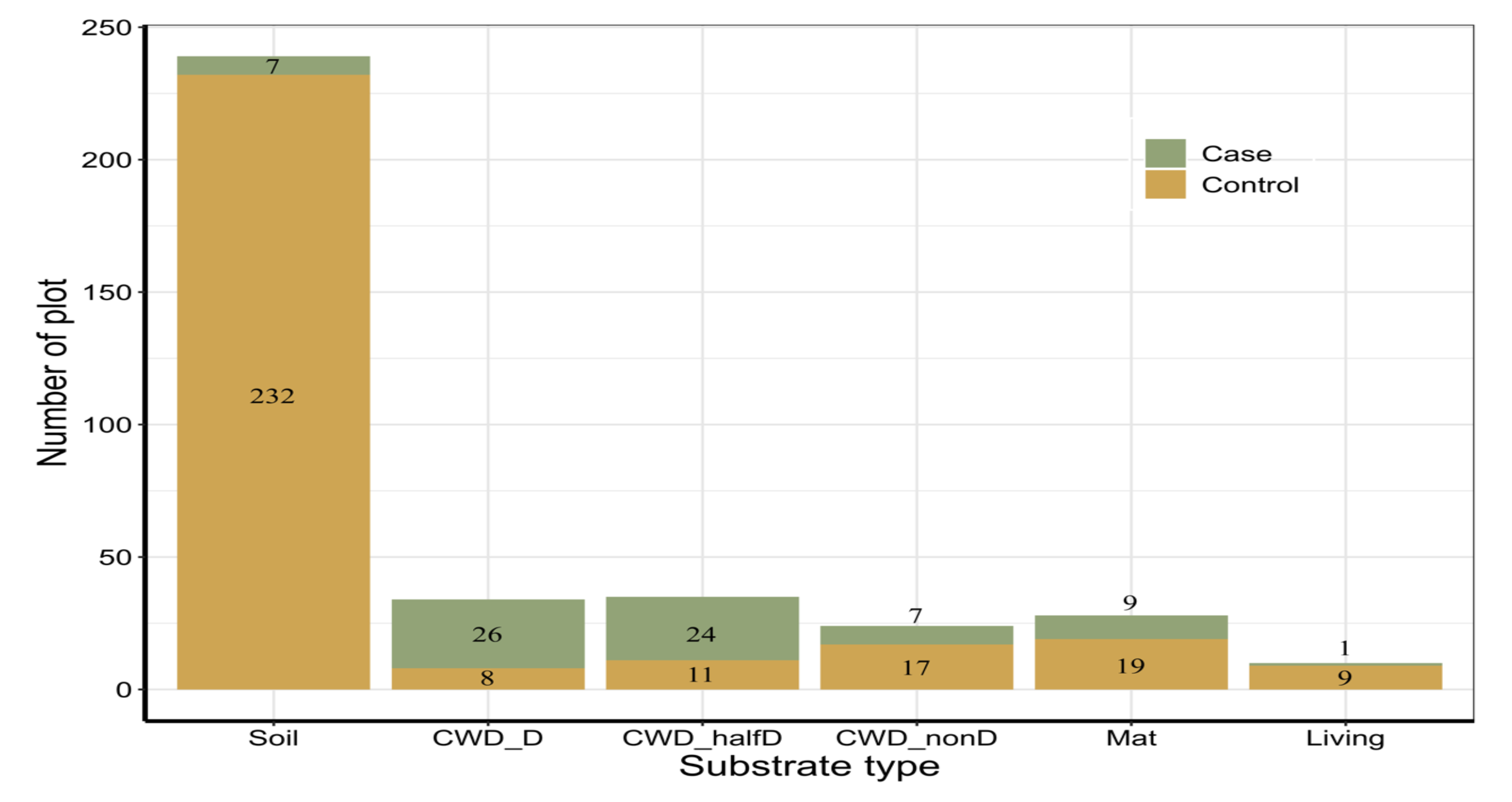
Numbers of plots on different substrate types. The numbers of plots were separated into case plots (green) and control plots (brown). The number on the bar indicates the actual numbers of the plot. (Soil represented forest soil; CWD_D, CWD_halfD, and CWD_nonD represented heavily-decayed, moderately-decayed, and non-decayed CWD, respectively; Mat represented organic matter mat; and Living represented living tree.)

Since there was not sufficient variance among regions, the mixed-effects conditional logistic regression model omitted the random intercept and was simplified as the ordinary conditional logistic regression model. The model included substrate types, bryophyte cover and thickness, litter thickness, and canopy cover as explanatory variables (Table 2). Bryophyte cover (Odds ratio (OR) = 7.654, p = 0.003) was significantly positively correlated with seedling occurrence. On the other hand, litter thickness (OR = 0.125, p = 0.017), bryophyte thickness (OR = 0.252, p = 0.044), and canopy cover (OR = 0.277, p = 0.015) were negatively associated with seedling occurrence. As for the substrate types, the model revealed that the effect of heavily-decayed CWD (OR = 24.328, p = 0.023) and moderately-decayed CWD (OR = 17.945, p = 0.016) were significantly superior to other substrates. The effect of living tree (OR = 0.498, p = 0.672) and non-decayed CWD (OR = 0.356, p = 0.419) were similar to soil. Mat (OR = 2.124, p = 0.522) had intermediate effect that was better than non-decayed CWD and soil but worse than heavy-decayed CWD. The adjusted McFadden’s pseudo-R^2^ was 77.3% (df = 9) and the concordance was 0.976 (standard error (SE) = 0.012). We further inspected whether mean fog frequency or precipitation would interact with substrate types. The model could not estimate the effect of the climatic variables since there were not enough between region variation.

**Table 2.** The result of conditional logistic regression of seedling establishment from case-control sampling. OR refers to the slope of the model (in logarithm) and should be interpret as odds ratio. Standard error (SE) is expressed as the log scale while confidence interval (CI) is in original scale. z refers to the z-statistic of the estimated parameters. Variables significant at p ≤ 0.05 are marked in bold font.

| Explanatory variables | log(OR) | OR | SE | z | p | CI |
| --- | --- | --- | --- | --- | --- | --- |
| Heavily-decayed CWD | 3.192 | 24.328 | 1.406 | 2.270 | **0.023** | 1.547 382.512 |
| Moderately-decayed CWD | 2.887 | 17.945 | 1.202 | 2.403 | **0.016** | 1.703 189.110 |
| Non-decayed CWD | -1.034 | 0.356 | 1.278 | -0.809 | 0.419 | 0.029 4.352 |
| Mat | 0.753 | 2.124 | 1.177 | 0.640 | 0.522 | 0.212 21.336 |
| Living tree | -0.697 | 0.498 | 1.644 | -0.424 | 0.672 | 0.020 12.508 |
| Bryophyte thickness | -1.378 | 0.252 | 0.685 | -2.011 | **0.044** | 0.066 0.966 |
| Litter thickness | -2.079 | 0.125 | 0.870 | -2.389 | **0.017** | 0.023 0.689 |
| Bryophyte cover | 2.035 | 7.654 | 0.681 | 2.987 | **0.003** | 2.013 29.099 |
| Canopy cover | -1.284 | 0.277 | 0.527 | -2.435 | **0.015** | 0.099 0.778 |

### Seedling survival monitoring

In the first survey in the summer of 2021, we labelled 869 seedlings in 36 plots across nine regions. The mean seedlings number per plot was 24 (ranging between 5 and 114). After four seasons of resurveys, 102 (11.7%) seedlings were considered to die as a result of environmental-induced mortality, 50 (5.8%) seedlings were marked as dead because of herbivory, and 81 (9.3%) seedlings went missing. The overall mortality rate was 26.8%. While from spring to summer, the death from environmental-induced mortality was larger than death from herbivory, from winter to spring, most of the deaths resulted from herbivory (Fig. 3). To unveil whether the seasonal differences of the causes of death were related to seasonal climatic variables, we firstly examined the effect of mean seasonal fog frequency, precipitation and temperature on seedling survival. For environment-induced mortality, the random effects of the model showed that regional variance was low (standard deviation (SD) = 0.020) compared to plot variance (SD = 0.658). None of the three climatic variables showed significant results (for mean seasonal fog frequency: HR = 1.129, p = 0.64; for mean seasonal precipitation: HR = −0.060, p = 0.94; for mean seasonal temperature: HR = 1.325, p = 0.81, Table A.2). For herbivory-induced mortality, similar results were found (for mean seasonal fog frequency: HR = 1.401, p = 0.33; for mean seasonal precipitation: HR = 13.381, p = 0.21; for mean seasonal temperature: HR = 0.590, p = 0.85, Table A.2). However, regional variance was relatively large (SD = 0.767) compared to plot variance (SD = 0.914). We further examined which climatic variables had enough differences to capture the heterogeneity of regional climatic characteristics (Table A.3). We found that mean annual precipitation significantly increased the hazard of the seedling being browsed (HR = 3.149, p < 0.001) while mean annual fog frequency and temperature were both insignificant (HR = 1.020, p = 0.940; HR = 1.442, p = 0.290).

**Fig. 3.**
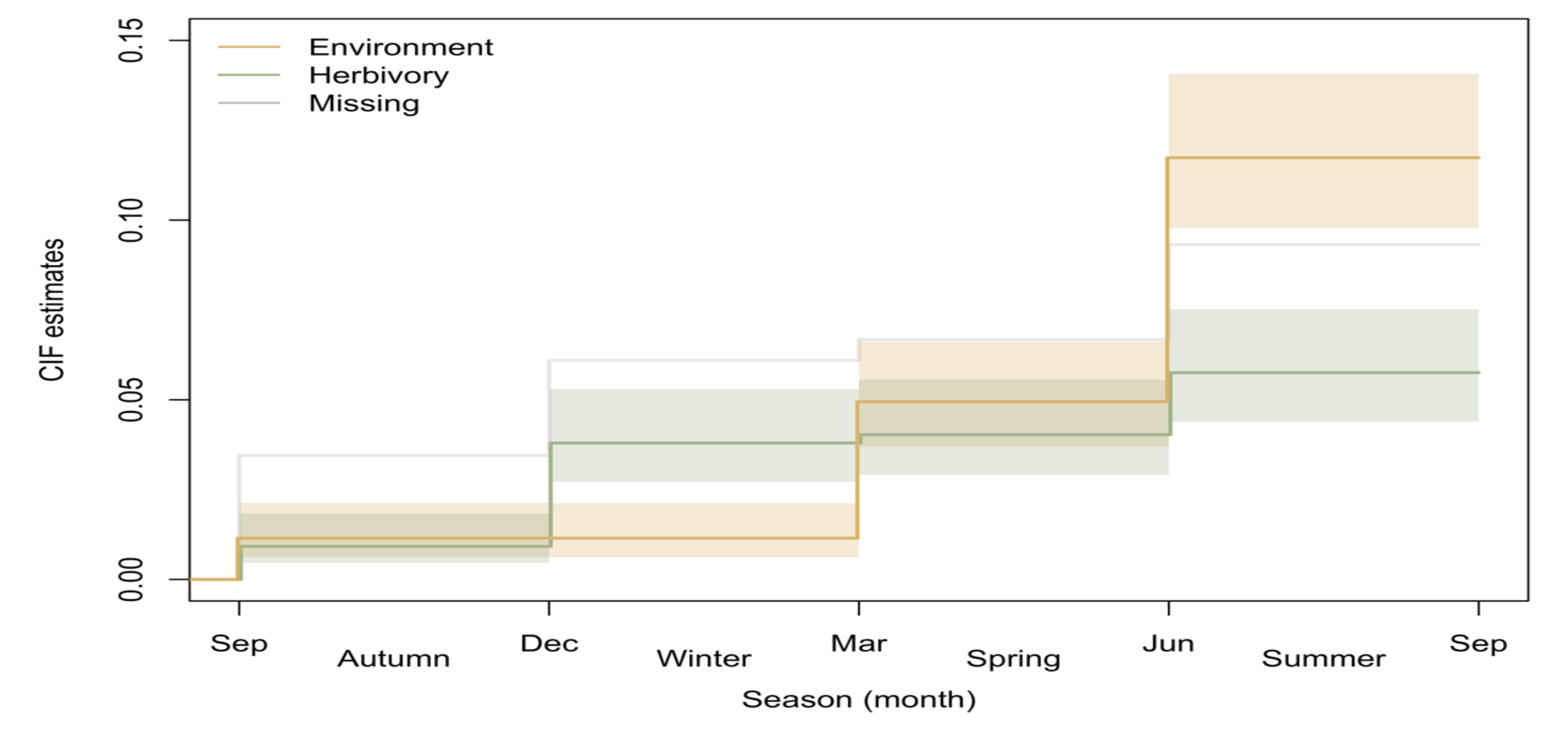
The cumulative incidence function (CIF) estimates for the competing risks of environmental-induced mortality or herbivory in the seedling survival monitoring. Brown line represented the mortality from environmental-induced mortality, green line represented the mortality from herbivory, and the grey line represented the missing individuals. The bands around the lines are calculated from Aalen-Johansen estimator to represent pointwise 95% confidence interval.

**Table A.2.** The results of cause-specific Cox proportional hazard (PH) models from seedling survival monitoring on regional time-dependent climatic variables. There are two competing risk factors, one is the mortality from environmental-induced mortality, and the other is the mortality from herbivory. HR refers to the slope of the model (in logarithm) and should be interpreted as hazard ratio (HR). The standard error (SE) is expressed as the log scale. Variables which have p ≤ 0.05 are marked in bold font. Random effect composed of two levels of nested random effect structure (plot nested in region) and expressed as standard deviation (SD).

| <u>Risk factor = Environmental-induced mortality</u> |  |  |  |  |  | <u>Risk factor = Herbivory</u> |  |  |  |  |  |
| --- | --- | --- | --- | --- | --- | --- | --- | --- | --- | --- | --- |
| (death = 102, chi-square = 10.64 (df = 18.63), $p = 0.059$) | | | | | | (death = 50, chi-square = 47.10 (df = 16.31), $p < 0.001$) | | | | | |
| Random effect: Region = 0.020, Region/Plot = 0.658 |  |  |  |  |  | Random effect: Region = 0.767, Region/Plot = 0.914 |  |  |  |  |  |
| Explanatory variables | log(HR) | HR | SE | z | p | Explanatory variables | log(HR) | HR | SE | z | p |
| Mean seasonal fog frequency | 0.121 | 1.129 | 0.259 | 0.47 | 0.64 | Mean seasonal fog frequency | 0.338 | 1.401 | 0.348 | 0.97 | 0.33 |
| Mean seasonal precipitation | -0.060 | 0.942 | 0.735 | -0.08 | 0.94 | Mean seasonal precipitation | 2.594 | 13.381 | 2.087 | 1.24 | 0.21 |
| Mean seasonal temperature | 0.281 | 1.325 | 1.151 | 0.24 | 0.81 | Mean seasonal temperature | -0.528 | 0.590 | 2.750 | -0.19 | 0.85 |

**Table A.3.** The results of cause-specific Cox proportional hazard (PH) models from survival monitoring on regional climatic characteristic variables. There are two competing risk factors, one is the mortality from environmental-induced mortality, and the other is the mortality from herbivory. HR refers to the slope of the model (in logarithm) and should be interpreted as hazard ratio (HR). The standard error (SE) is expressed as the log scale. Variables which have p ≤ 0.05 are marked in bold font. Random effect composed of two levels of nested random effect structure (plot nested in region) and expressed as standard deviation (SD).

| <u>Risk factor = Environmental-induced mortality</u> |  |  |  |  |  | <u>Risk factor = Herbivory</u> |  |  |  |  |  |
| --- | --- | --- | --- | --- | --- | --- | --- | --- | --- | --- | --- |
| (death = 102, chi-square = 13.63 (df = 17.16), $p = 0.018$) | | | | | | (death = 50, chi-square = 57.75 (df = 11.87), $p < 0.001$) | | | | | |
| Random effect: Region = 0.020, Region/Plot = 0.623 |  |  |  |  |  | Random effect: Region = 0.020, Region/Plot = 0.789 |  |  |  |  |  |
| Explanatory variables | log(HR) | HR | SE | z | p | Explanatory variables | log(HR) | HR | SE | z | p |
| Mean annual fog frequency | 0.198 | 1.220 | 0.179 | 1.11 | 0.27 | Mean annual fog frequency | 0.020 | 1.020 | 0.264 | 0.08 | 0.940 |
| Mean annual precipitation | 0.029 | 1.029 | 0.188 | 0.15 | 0.88 | Mean annual precipitation | 1.147 | 3.149 | 0.304 | 3.77 | **<0.001** |
| Mean annual temperature | 0.131 | 1.140 | 0.165 | 0.80 | 0.43 | Mean annual temperature | 0.366 | 1.442 | 0.349 | 1.05 | 0.290 |

Individual-and plot-levels factors were also explored to quantify their association with seedling survival. Cause-specific Cox PH model have shown that deaths from environmental-induced mortality and herbivory were associated with different abiotic and biotic factors at local scale. For the environment-induced mortality (Table 3), variables selected in the final model included litter covered, bryophyte thickness and cover, and herb cover. The hazard ratio (HR) increased by 3.955 times (p < 0.001) for the seedlings being covered by litterfall. Bryophyte thickness (HR = 0.143, p < 0.001) and bryophyte cover (HR = 0.679, p = 0.010) were both shown to reduce the hazard of death of the seedlings. Interestingly, the effect of the bryophyte thickness in reducing the HR decreased linearly (HR = 1.625, p = 0.003) over time. Herb cover (HR = 11.756, p < 0.001) would increase the hazard, but its effect (HR = 0.537, p < 0.001) reduced over time. The concordance of the model was 0.645 (SE = 0.041) with pseudo-R^2^ = 0.392. For herbivory-induced mortality, the final model included bryophyte thickness, herb cover, litter cover, canopy cover, seedling height, and distance to the soil surface as explanatory variables (Table 3). Seedling height (HR = 1.776, p < 0.001) and bryophyte thickness (HR = 1.960, p = 0.002) both significantly increased the hazard of the seedlings being browsed. Herb cover (HR = 0.433, p = 0.003), distance to the soil surface (HR = 0.570, p = 0.018), and canopy cover (HR = 0.404, p < 0.001) reduced the hazard and did not vary over time. The effect of litter cover was insignificantly (HR = 0.807, p = 0.662) associated with the hazard in the initial state but reduced the risk linearly over time (HR = 0.660, p = 0.023). The concordance was 0.767 (SE = 0.062) and pseudo-R^2^ = 0.613 for the model.

**Table 3.** The results of cause-specific Cox proportion hazard (PH) models from seedling survival monitoring. There are two competing risk factors, one is the mortality from environmental-induced mortality, and the other is the mortality from herbivory. HR refers to the slope of the model (in logarithm) and should be interpreted as hazard ratio (HR). The standard error (SE) is expressed as the log scale. Variables within tt() refer to the time-varying variables for which the hazard ratio changes over time. Variables which have p ≤ 0.05 are marked in bold font.

| <u>Risk factor = Environmental-induced mortality</u> |  |  |  |  |  | <u>Risk factor = Herbivory</u> |  |  |  |  |  |
| --- | --- | --- | --- | --- | --- | --- | --- | --- | --- | --- | --- |
| (death = 102, Wald-statistic = 66.11, $p < 0.001$, Concordance = $0.645 \pm 0.041$) | | | | | | (death = 50, Wald-statistic = 69.62, $p < 0.001$, Concordance = $0.767 \pm 0.062$) | | | | | |
| Explanatory variables | log(HR) | HR | SE | z | p | Explanatory variables | log(HR) | HR | SE | z | p |
| Litter covered | 1.375 | 3.955 | 0.301 | 4.760 | **<0.001** | Distance to the soil surface | -0.562 | 0.570 | 0.258 | -2.375 | **0.018** |
| Bryophyte thickness | -1.944 | 0.143 | 0.649 | -3.269 | **0.001** | Seedling height | 0.574 | 1.776 | 0.152 | 5.410 | **<0.001** |
| Bryophyte cover | -0.387 | 0.679 | 0.154 | -2.592 | **0.010** | Canopy cover | -0.907 | 0.404 | 0.395 | -3.437 | **0.001** |
| Herb cover | 2.464 | 11.756 | 0.795 | 4.312 | **<0.001** | Bryophyte thickness | 0.673 | 1.960 | 0.283 | 3.027 | **0.002** |
| tt(Bryophyte thickness) | 0.486 | 1.625 | 0.178 | 2.934 | **0.003** | Litter cover | -0.215 | 0.807 | 0.679 | -0.437 | 0.662 |
| tt(Herb cover) | -0.621 | 0.537 | 0.218 | -4.103 | **<0.001** | Herb cover | -0.837 | 0.433 | 0.420 | -2.924 | **0.003** |
|  |  |  |  |  |  | tt(Litter cover)) | -0.415 | 0.660 | 0.210 | -2.268 | **0.023** |

## Discussion

Our study applied two sampling designs and analysis techniques to identify the stress factors encountered by the *Chamaecyparis obtusa* var. *formosana* seedlings, and to quantify the impact of regional climatic variables and local biotic and abiotic factors in forming the seedlings’ safe site in terms of establishment and survival. The findings were interconnected since the combination of establishment and survival determined the seedlings’ final success.

After monitoring the seedling survival seasonally for a year, we identified two alternative ways of seedling death: environmental-induced mortality or herbivory. For environmental-induced mortality, although we set the sampling regions along a gradient of fog frequency to capture various climatic characteristics, we found that different sampling regions each encompassed similar seasonal trends without strong regional variation. Approximately 12% of seedlings died from environmental-induced mortality, especially in spring and summer. This may result from the rainfall gap between the Mei-Yu (Plum rain) and typhoon seasons, during which the evapotranspiration rate usually increases as temperature increases and fog frequency declines (Chen, 1994; Chen and Chen, 2003; Chang et al., 2006; Lai et al., 2006; Chu et al., 2014). The rainfall gap creates short-duration drought that could be stressful to the cloud forest species; therefore, we suspect this phenomenon might be the underlying mechanism of the environment-induced mortality (Liao et al., 2003a; Chu et al., 2014; Oliveira et al., 2014). On the other hand, no environmental-induced mortality was observed during the winter, suggesting that low-temperature stress, including frost events, may not be the limiting factor. Unfortunately, we do not have enough evidence to directly confirm which climatic factor might be the underlying mechanism by analyzing the relationship between seasonal climatic variables to the survival of the seedlings. This may be because the time resolution in our climatic data was too coarse (averaging across three months) to capture the short-period drought event or the strong inter-annual fluctuation during this period might be smoothened out when averaging the data across different years (Wu et al., 2018).

In addition to environmental-induced mortality, we also observed that 6% of seedlings died due to browsing by herbivores. This death rate showed substantial regional variation along a gradient of annual precipitation, with higher rainfall regions having more dead seedlings due to herbivory. For instance, in the Lalashan and Tamanshan regions, over 15% of individuals died from herbivory, while in Daguangshan, it was 39% of seedlings. The two main herbivores in the mid-elevation of Taiwan are barking deer (*Muntiacus reevesi micrurus*) and Formosan serow (*Capricornis crispus swinhoei*). Both species prefer gaps in the forest and are generalists with a diet mainly composed of tender leaves and fruits from herbs (e.g., *Rubus* spp.) and shrubs, and usually suffer from food shortages in winter (Lue, 1987; McCullough et al., 2000). It is therefore not surprising that a closed canopy and higher litter cover reduced the herbivory stress on seedlings in our data. However, we still cannot satisfactorily explain the regional correlation between herbivory pressure and mean annual precipitation.

Under the stresses of either environmental-induced mortality or herbivory, our analysis suggests that the safe site requirements of the seedlings need to balance between minimizing the negative effects from environmental-induced mortality and herbivory to maximize the seedlings’ survival. For example, under dense herb cover, seedlings may experience strong light and nutrients limitation, but the same herb cover can also act as a shelter, reducing herbivory pressure. Similarly, litter accumulation threatened the seedlings by overtopping them, but sites with higher litter cover might attract fewer herbivore visits, thereby reducing browsing pressure. Our study highlights the complex interplay between different stress factors on seedling survival. Ignoring one of these stress factors can lead to a less precise or incorrect interpretation of the ecological significance of the associated variables.

In light of the complex interplay between environmental-induced mortality and herbivory on seedling survival and establishment, the overall impacts of the abiotic and biotic factors on the safe site requirements of the seedlings were explored by the seedling establishment survey supplemented with information from survival monitoring. At the regional scale, we found that regional climatic characteristics, specifically fog frequency and precipitation gradient, exhibited minimal variance across different regions and had no discernible impact on seedling establishment and survival. Notwithstanding the lack of evidence in results of our study, we did observe that the seedlings appeared to be less confined to decayed CWD (including moderately-decayed and heavily-decayed CWD) in moist regions (such as Donqiusishan). This observation suggests that the regional climatic characteristics do affect seedlings’ safe site requirements, at least with respect to substrate types.

Contrary to regional climatic variables, local-scale factors, such as substrate types, bryophyte thickness and cover, litter thickness, and canopy cover, were crucial in determining whether the seedlings could establish; this however did not apply to the effect of heterospecific competition. Except for canopy cover, most of crucial local-scale factors are intercorrelated. In particular, different substrate types have distinct associations with varying cover and thickness of bryophytes and litter. Decayed CWD is associated with a higher cover of small-stature bryophytes and lower litter cover, while soil has the opposite pattern with large-stature bryophytes or a higher and thicker litter cover. Despite these relationships, decayed CWD still has a significant positive effect on seedlings’ establishment when all variables were analyzed together, without strong multicollinearity. This suggests that there might be intrinsic properties of decayed CWD that better support seedling establishment, compared to non-decayed CWD and soil. From the abiotic perspective, decayed CWD has a greater water-holding capacity which reduces moisture fluctuation under drought stress (Pitchler et al., 2011), and also lower nutrient status compared to soil (Takahashi et al., 2000; Sakai et al., 2012; Wei and Zelený, 2023). Also, we observed that most of the decayed CWD in the field were affected by brown-rot fungi which would release more sugar for fungi growth and create more crevice for seedling rooting (Wang et al., 1980; Chang and Duh, 1988; Fukasawa, 2018). On the other hand, decayed CWD might reduce the competition pressure since other species might not be able to adapt to low nutrient status and the allelopathic effects resulting from high secondary compounds concentration on *Chamaecyparis* wood (Chang and Duh, 1988; Harmon and Franklin, 1989; Tseng et al., 2007; Sakai et al., 2012). Furthermore, the position of decayed CWD elevated above the surrounding substrate may act as a physical barrier for the herbivores, thereby reducing the risk of browsing (Hagge et al., 2019).

In addition to the role played by the substrates, our results also indicated that various bryophyte carpets covering the top of different substrates might have contrasting effects on the seedling establishment. Large-stature and thick bryophyte cover, which is more frequently found on forest soils, negatively affected the seedling’s establishment by competing for water during drought period, interfering with seedlings to reach the beneath humus layer, or even attracting more herbivores (Nakamura, 1992). We speculate that compared to the thick tall turf bryophyte species like *Polytrichum* or *Sphagnum*, the thinner short turf or wefts bryophyte community associated with decayed-CWD (e.g., *Bazzania, Pyrrhobryum*, and Sematophyllaceae species) are particularly beneficial to the seedlings (Fukasawa and Ando, 2018). Several studies also suggested that thin and small-statured bryophyte cover might facilitate seedling establishment via humus accumulation and thus have better moisture and nutrient regimes without strong competition (Harmon et al., 1986; Takahashi, 2000; Iijima et al., 2006; Wei and Zelený, 2023). Bryophytes have also been reported to intercept and store nutrients and water from rainfall or fog and subsequently act as slow-release fertilizer, especially under drying events (Clark et al., 2005; Ah-Peng et al., 2017; Glime, 2017). Additionally, bryophyte cover might trap more seeds from the seed rain than the bare CWD surface to germinate, especially for small-seed species like *Chamaecyparis obtusa* var. *formosana* (Harmon and Franklin, 1989). Aside from the role of bryophyte community, our study did not take into account the potential role of fungi, particularly saprotrophic and mycorrhiza fungi inhabiting on CWD, which might ameliorate the low-nutrient situation by decomposition and determine the bryophytic community composition (Fukasawa et al., 2015; Fukasawa et al., 2017). Future studies could explore the relationship between fungi community and CWD decomposition, bryophyte community structure and seedling establishment.

In contrast to the effect of bryophyte carpets, which varied according to the bryophyte stature, litter dynamics had a constantly negative impact on seedling persistence. A thick and dynamic litter layer, which usually accumulates on the forest soil surface, might hinder the seeds from reaching the soil for germination (Wang et al., 2022) or prevent seedling emergence (Knapp and Smith, 1982; Simard et al., 2003). Gillman (2016) pointed out that the combination of macrolitterfall amount and seedling resilience influenced the damage of the seedlings caused by litterfall. From personal observation, we can see that litterfall was composed mainly of the twigs and leaves of *Chamaecyparis obtusa* var. *formosana* and tough leaves of Fagaceae or Rhododendron spp. (Lu and Liu, 2012). The litter is usually larger and heavier than the seedlings, especially for those seedlings which haven’t developed harden stem or haven’t reached enough height. When covering by the hard-to-decompose litters, we noticed many of the seedlings eventually died.

Beyond the factors associated with substrate types, differences in light regimes also played an important role in successfully establishing *Chamaecyparis obtusa* var. *formosana* seedlings. Previous studies have demonstrated that these seedlings exhibit better growth under moderate light intensity, with the ability to adapt quickly to sunflecks (Fang et al., 1991; Lai et al., 2005). Conversely, high light intensity can lead to a reduction in the relative growth of *Chamaecyparis* leaves. Our analysis found a linear negative correlation between the estimated canopy cover and seedling establishment. This could be because we only considered light regimes ranging from low to moderate light conditions in closed mature forests, which the seedlings can tolerate and thrive in, while full light conditions, which inhibit their growth under disturbed forests, were not included in our study. Additionally, the understory herb layer may have altered the light regime, with higher herb cover reducing light availability for the seedlings and thus possibly having the adverse effect on individual survival. However, since the total cover of the herb layer is small, the negative effect is negligible at whole plot-scale.

From the analysis of seedlings’ survival and establishment data, we have rather limited evidence that regional climatic variables would impact the microhabitat requirements and cause stress amelioration of the seedlings. On the other hand, we did observe a close relationship between decayed CWD and bryophyte cover, which might play a crucial role in mitigating the short-duration drought stress during spring and summer seasons to facilitate seedling establishment process. Such relationship between the high water-holding capacity of CWD and bryophyte carpet to ameliorate drought of seedlings was also documented in other small-seed conifer species worldwide. For example, it has been shown that *Picea sitchensis* (Bong.) Carr. and *Tsuga heterophylla* (Raf.) Sarg. in the Pacific Northwest region favor CWD, benefiting from their higher moisture content when facing summer drought (Gray and Spies, 1997). In Japan, elevated decayed CWD is a better regeneration site for seedlings of *Chamacyparis obtusa* (Sieb. and Zucc.) Endl.*, Picea glehinii* (Fr. Schm.) Masters, and *P. jezoensis* (Sieb. et Zucc.) Carr., since CWD and associated bryophyte carpets have a better water-holding capacity that meets the requirement of the seedling establishment (Yamamoto, 1993; Takahashi et al., 2000; Fukasawa and Ando, 2018). In our study, we hypothesize that the short-duration drought might be a potential drought stress factor. Studies predicting future climate indicate that the prolonged dry days and drought, which increased since 1960, will continue to increase in the future climate change scenario (Cheng et al., 2009; Huang et al., 2012). Other regions may also experience similar drought stresses due to various local climatic phenomena. As droughts intensify due to climate change (Huang et al., 2012), the seedlings might be more confined to decayed CWD with bryophyte cover to resist drought stress. To sustain the population of species that rely on decayed CWD and the associated bryophyte community for regeneration, it is crucial to retain CWD in the forest floor for natural degradation after tree-replacing disturbance in natural forests.

## Conclusions

In our study, we integrated seedling survival monitoring and establishment survey to identify the stress factors and reveal the safe site requirements for the early-stage establishment of *Chamaecyparis obtusa* var. *formosana* seedlings. Across the species’ northern distribution range in Taiwan under different climatic conditions, we identified that environmental-induced mortality and herbivory were the possible alternative ways of seedling death. Interestingly, we also found that seedlings might need to balance between favorable ecological factors under both stressors to optimize their survival and establishment. Therefore, we propose that the decayed CWD serves as a “safe site” for the seedlings, regardless of climatic regimes. However, the underlying mechanism might not only be the decayed CWD itself but also because it nurtures the bryophyte cover supporting the seedlings’ growth and avoids the damage to seedlings caused by litterfall. Overall, our study highlights the complex interplay between different stress factors and local environmental variables and points out the need to conserve CWD after tree-replacing disturbances in natural forests to ensure the sustainability of population growth. However, we did not consider the potential role of fungi inhabiting on CWD. Future studies should explore the relationship between fungi community and CWD decomposition, bryophyte community structure, and seedling establishment. Additionally, further research is needed to track the seedling-to-sapling transition stage and quantify the environmental requirements in that stage to provide insights into the conservation and management of *Chamaecyparis obtusa* var. *formosana* populations in Taiwan’s forests.

## Acknowledgment

The authors sincerely thank Chi-Cheng Liao, Biing-Tzung Guan, Koh-Suan Chen, and Jéssica Viana for constructive comments. The authors also wish to acknowledge all the people who make invaluable contribution in the field during the development of this research, in particular to Ming-Wei Zhong, Yi-Mei Wu, Tsung-Chen Lee, Shu-I Lin, Hsun-Hung Chu, Yin-Sun Yao, Ching-Lin Huang, and Kun-Sung Wu. Funding was provided by the Ministry of Science and Technology, Taiwan (MOST, 109-2621-B-002- 002-MY3).

## Notes

https://github.com/zdealveindy/VegLab/tree/main/data/chamaecyparis-seedlings-establishment-survival

### Competing Interest Statement

The authors have declared no competing interest.

